# An Efficient Coalescent Epoch Model for Bayesian Phylogenetic Inference

**DOI:** 10.1101/2021.06.28.450225

**Authors:** Remco R. Bouckaert

**Affiliations:** University of Auckland, Auckland, New Zealand

## Abstract

We present a two headed approach called Bayesian Integrated Coalescent Epoch PlotS (BICEPS) for efficient inference of coalescent epoch models. Firstly, we integrate out population size parameters and secondly we introduce a set of more powerful Markov chain Monte Carlo (MCMC) proposals for flexing and stretching trees. Even though population sizes are integrated out and not explicitly sampled through MCMC, we are still able to generate samples from the population size posteriors. This allows demographic reconstruction through time and estimating the timing and magnitude of population bottlenecks and full population histories. Altogether, BICEPS can be considered a more muscular version of the popular Bayesian skyline model.

We demonstrate its power and correctness by a well calibrated simulation study. Furthermore, we demonstrate with an application to SARS-CoV-2 genomic data that some analyses that have trouble converging with the traditional Bayesian skyline prior and standard MCMC proposals can do well with the BICEPS approach.

BICEPS is available as open source package for BEAST 2 under GPL license and has a user friendly graphical user interface. Bayesian phylogenetics, coalescent model, BEAST 2, BICEPS

## 1 Introduction

Knowledge of population size dynamics can be of interest, for example, for the study of megafauna extinctions Campos et al. (2010), conservation biology (Shapiro et al., 2004), reconstructing human settlement history (Pedro et al., 2020), impact of viral ecology on public health (Rambaut et al., 2008), or the influence of climate events on population sizes Miller et al. (2012). Here, we will infer population size dynamics using a phylogeny with sequence data on a single gene, for example, mitochondrial sequences, or full genome viral data, based on coalescent theory in a Bayesian setting. We do not assume any structure, that is, we assume there is a single population, and we assume there is random mating and no admixture. Coalescent theory links phylogenies with population sizes through tree priors based on Kingman’s theory (Kingman, 1982). These tree priors are driven by a population function that define the effective population size through time. A population function can be parametric, like exponential or constant (Kuhner et al., 1998), but non-parametric methods that split up the time frame spanning a tree into epochs allow a population function to be constant in an epoch but vary over time. Non-parametric methods allow representation of a much wider range of population functions than parametric methods, and can capture one or more population bottlenecks and expansions without a priori having to commit to the number of such bottlenecks or expansions. So, nonparametric models offer a flexible alternative to parametric models and allow more wide range of population size dynamics estimates. Even when population size dynamics is of no interest, these models provide a flexible tree prior allowing a broad range of tree shapes and sizes.

The classic skyline model (Pybus et al., 2000), introduced in a maximum likelihood framework, is based on epochs for every coalescent event. It assumes that the phylogeny is fully resolved and divergence time estimates are reliable, so can only be applied when the data exhibits strong phylogenetic signal. The classic skyline model was later generalised to epochs grouping coalescent events in the generalised skyline model (Strimmer and Pybus, 2001), making it possible to estimate population histories when little divergence information is available, for instance, when the alignment contains identical sequences. The Bayesian skyline plot (Drummond et al., 2005) generalised this to a Bayesian setting, where epochs span multiple coalescent events, and the number of coalescent events as well as population sizes for an epoch are sampled during MCMC. Furthermore, a smoothing prior is employed that links population sizes in consecutive epochs. Linking population sizes reduces stochastic noise and make biological sense in that consecutive population sizes will usually be of a similar order of magnitude as preceding ones. Other popular epoch based coalescent models with different smoothing priors include the skyride prior (Minin et al., 2008), which takes the amount of time between epochs in account, the skygrid prior (Gill et al., 2013; Hill and Baele, 2019), which allows users to define epoch boundaries, and the besp model (Parag et al., 2020), which takes sampling times in account.

All the above Bayesian methods sample the population function parameters. By assuming an inverse gamma prior distribution on population size, we demonstrate that the population size can be integrated out during MCMC. The technique is used in the multispecies coalescent models StarBeast2 (Ogilvie et al., 2017) and STACEY (Jones, 2017), where a constant population size is associated with each branch of the species tree. Here, we generalise this method to the case where we have a single tree, potentially with sampled tip dates, and assume a piecewise constant population for each epoch under an inverse gamma prior. The mean for the population size of the youngest epoch can be sampled but for consecutive epochs the posterior mean of the previous epoch can be used, providing us with a smoothing prior. Even though population sizes are integrated out, they can still be sampled from the posterior population sizes for the epochs conditioned on the tree. This allows us to reconstruct population size history including uncertainty intervals in a similar fashion as for the Bayesian skyline plot as follows. At regular intervals during the MCMC we log the tree, group sizes and sample for each group a population size from the posterior. Each such sample defines a demographic history where the length of each epoch is defined by the tree and group sizes, a so called skyline plot (Fig 1 Drummond et al. (2005)). So, for each point in time the skyline plot defines a population size for a particular tree and its parameters. By considering all the trees and other parameters in the posterior we get a distribution of population sizes for each point in time, which we can use to find the confidence intervals of the distribution (Fig.5).

**Figure 1:**
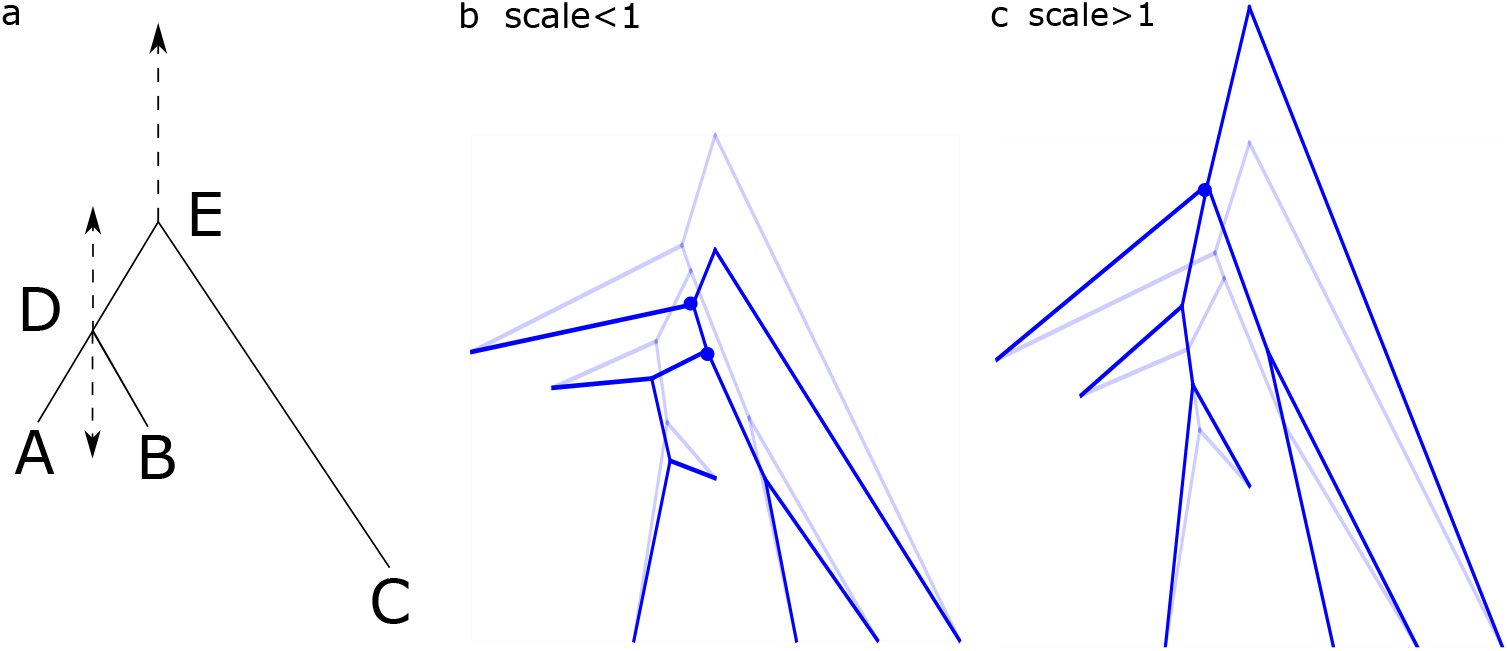
a) The traditional scale operator that gets often rejected when there are many tip dates (*r* ≫ 1) because of high probability of negative branch lengths when scaling down or inappropriately stretching short branches into older tips when scaling up. Tree stretch proposal moves nodes near tips (e.g. node D) less far than nodes away from tips (e.g. node E) b) for scale factor less than 1 where lighter trees are the original state and darker trees are proposals and c) for scale factor larger than one.

Apart from introducing a more efficient way to infer population size histories at different epochs, we also consider a number of new MCMC proposals that can lift a large number of nodes in a tree simultaneously. Observing that tree priors tend to be highly correlated with the length of a tree, we target tree length changes by moving nodes in randomly chosen time intervals (not necessarily the ones used for the tree prior). Note that we are considering rooted time trees only, so the tree length is defined as the sum of branch lengths in units of time of the tree. The likelihood is also correlated with the length of the tree, but only after scaling it with a clock rate.

Furthermore, noting that scaling of trees tends to be hampered by serially sampled tips we design a new scale proposal that moves all nodes in a tree simultaneously but with better exploratory powers than standard scalers. Both proposals move tree length, and since clock rates tend to be inversely correlated with the tree length (Douglas et al., 2021c), we designed proposals that simultaneously move the clock rate to compensate for a changing tree length. We demonstrate the effectiveness of these MCMC proposals for improving mixing of tree lengths, and thus tree priors.

Together, integrating out population sizes and employing more sophisticated MCMC proposals allow us to do inference efficiently, and make it possible to perform larger analyses, as we demonstrate using SARS-CoV-2 data. In the next section, we consider the technical details around integrating out parameters and new MCMC proposals. We continue with validating the method and presenting results. In the conclusions (final section) we consider ways to generalise the approach and in particular point out how to integrate out parameters for an epoch version of the Yule prior (Appendix B and C).

## 2 Methods

First, we consider integrating out population size parameters, then we design a set of new MCMC proposals.

### 2.1 BICEPS model: integrating out parameters

Let *T* be a rooted binary tree with *n* taxa sampled at *r* different times.^1^ So, *r* = 1 when all taxa are sampled at the same time and *r* = *n* when all taxa are sampled at different times. Then there are *n* + *r* − 2 times *t*_1_, *t*_2_, …, *t*_*n*+*r−*2_ that are either sampling times or coalescent times ordered from youngest tip (*t*_1_) to the the coalescent time at the root (*t*_*n*+*r−*2_) and let Δ*t_i_* = *t_i_* − *t*_*i−*1_ denote the length of an interval. Let *k_i_* be the number of lineages at event *i*, so *i* decreases by one at a coalescent event, but increases at a sampling event. Let 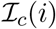 be an indicator function that indicates whether the *i*th event is a coalescent event 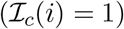 or a sampling event 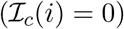.

Consider *m* epochs defined by groups of coalescent events, and let *A* = {*a*_1_, *a*_2_, …, *a_m_*} be the number of coalescent events in each of the *m* epochs that cover the whole tree (so 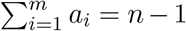). Parag and Pybus (2019) show that having a similar number of coalescent events per epoch increases accuracy of population size estimates, so in practice we keep group sizes constant and evenly spread. The number of epochs is a parameter to be provided by the user, but by default 10 epochs will be used unless the epoch sizes become less than 6 (*n/*6 groups will be used) or larger than 30 (*n/*30 groups will be used).

Let Θ = {*θ*_1_, *θ*_2_, *…, θ_m_*} be the effective population sizes for the *m* epochs, that define a piecewise constant population function for the *m* epochs. Let *h*(*i*) be a function 1, *…, n*+*s*−2 → *m* that map the coalescent and sampling events *i* to epochs (Drummond et al. (2005) Eq(4)). Then the log likelihood log *p*(*T* |Θ, *A*) of the tree *T* given Θ and *A* is (Drummond et al. (2005) Eq(3)):

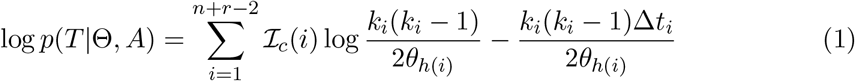

Taking the exponent, gives the density

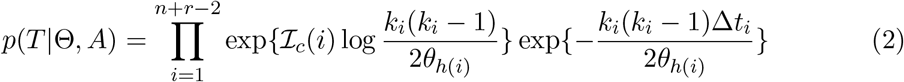

Let *p_j_*(*T* |Θ, *A*) denote the contribution for a single epoch *j* so 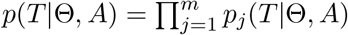, and let *b_j_* be the index of event *i* at the start of the *j*th epoch (so, *h*(*i*) = *j* for *b_i_* ≤ *i* < *b_i_*_+1_), then

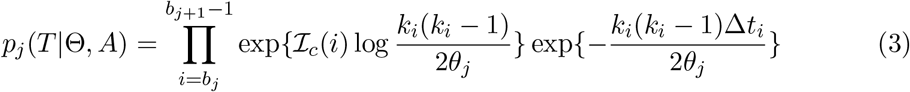

which can be simplified to

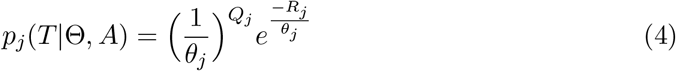

with 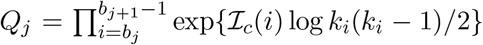 and 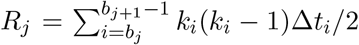. Following Liu et al. (2008), we note that the inverse gamma distribution 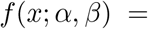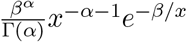 is conjugate for *θ_j_*, in other words, the posterior is *f* (*θ_j_*|*α* + *Q_j_, β* + *R_j_*) and integrating out *θ_j_* gives

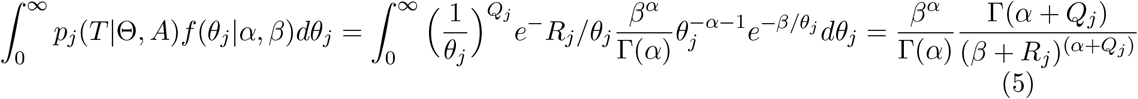

thus we get a closed form density for the contribution of epoch *j* that has the population size *θ_j_* integrated out. Since 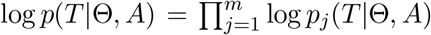 and *θ_j_* independent, we can do this for each of the intervals.

This leaves us to choose the parameters for the inverse gamma prior on population sizes. If no further information about population sizes, this prior ideally has little influence on the distribution of population sizes (Liu et al., 2008). By default, the shape value of *α* = 3 is fixed as suggested elsewhere (Ogilvie et al., 2017; Liu et al., 2008), which has the special property that the standard deviation is identical to the mean (Ogilvie et al., 2017), so the coefficient of variation is 1, providing a wide ranging distribution. If there is some information about possible values of *α* these can be changed. The population mean 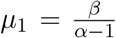 estimated during the MCMC run with a lognormal(*μ* = 1, *σ* = 1) by default.

#### 2.1.1 Smoothing priors

Epoch models can show abrupt changes in population size estimates when population sizes for the epochs are assumed to be independent. For that reason, smoothing priors are applied (Drummond et al., 2005; Minin et al., 2008; Gill et al., 2013), which suppress large fluctuations of population sizes in consecutive epochs. One way to do this is to sample only the population mean for the first epoch, and for consecutive epochs, the posterior mean of the previous epoch 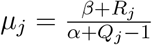 can be used to set *β*_*j*+1_ = *μ*_*j*_ (*α* - 1).

#### 2.1.2 Inferring skyline plots

While models that explicitly sample population sizes of each epoch store population sizes and epoch information during MCMC, we do not have population size information available when integrating them out. However, given that for each epoch *j* we have a posterior distribution *f* (*θ_j_*|*α*+*Q_j_, β_j_* +*R_j_*) we can just sample a value from that posterior and approximate the population size distribution for each epoch, and this allows us to perform demographic reconstruction. A sample from an inverse gamma distribution can be obtained by sampling a gamma distribution with shape *α* + *Q_j_* and scale 1*/*(*β_j_* + *R_j_*) and taking the reciprocal value of the sample.

### 2.2 BICEPS operators

To help convergence of the MCMC algorithm, we introduce a number of new proposals that move a large number of heights of internal nodes in the tree while keeping leaf node heights constant. These proposals have a large effect on the length of a tree, and thus indirectly on the tree prior. Note that the methods introduced are applicable to all phylogenetic tree priors and are not restricted to the epoch model discussed above.

#### 2.2.1 New tree stretch proposal

The standard tree scale proposal in BEAST 2 simply multiplies all internal node heights *h_i_* (for node *i*) with the same randomly chosen scale factor *s*, but leaf node heights remain unchanged. This can lead to negative branch lengths if an internal node is scaled down below a tip height of a descendant, at which point the scale proposal is instantly rejected (see node D in Fig. 1a when scaled down). When there are many dated tips over a large time range, and there is little variation in sequence data resulting in short terminal branches, scaling up can make relatively short terminal branches stretch out a lot causing a marked reduction in tree likelihood causing the proposal to be rejected (see node D in Fig. 1a when scaled up).

To remedy such low acceptance, the range from which the scale factor is sampled can be reduced, but that leads to smaller overall changes to the tree. Note that when scaling all nodes in the tree the pruning algorithm (Felsenstein, 1981) for calculating the tree likelihood needs to recalculate all so called partials for internal nodes, which is a computationally expensive task (see (Felsenstein, 1981) for details). So, ideally we would like to make bold proposals to justify this computationally costly operation.

Instead of simply multiplying internal node heights, as the standard scale operator does, we can do a post-order traversal where we scale branch lengths and add them to the height of the left and right child, then take the average of these heights to set the height of the current node in the traversal. Formally, let *s* be a randomly chosen scale factor from a Bactrian kernel (Yang and Rodríguez, 2013; Thawornwattana et al., 2018), that is, we randomly sample a value from a standard Gaussian *N* (0, 1) scaled with 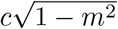, and randomly add or subtract *m*. Here, *m* determines the shape of the Bactrian distribution and is set to 0.95 by default, and *c* is a tuning parameter. The tuning parameter is automatically optimised (Drummond and Bouckaert, 2015) during MCMC to obtain optimal balance between better acceptance (at lower values of *c*) and boldness (at larger values of *c*). A target acceptance probability of 0.4 suggested in Yang and Rodríguez (2013) appears to give good results. Automatic tuning of operators ensures that for models with high rejection rate, the size of the proposed changes will be reduced, so subsequent proposals will be less bold. Let *b_i_* be the branch length above node *i*, so *b_i_* = *h_p_* − *h_i_* when *p* is the parent of node *i*. We traverse the tree and do not change leaf node heights, but for a node *i* with children *j* and *k* (assuming they were already visited), we set the new height *h′_i_* of node *i* to (*h′_j_* + *sb_j_* + *h′_k_* + *sb_k_*)*/*2. When all tips are contemporary, this proposal is the same as the traditional tree scale operator (because *h′_i_* = (*h′_j_* +*sb_j_* + *h′_k_* +*sb_k_*)*/*2 = (*h′_j_* + *s*(*h_i_*−*h_j_*)+ *h′_k_* + *s*(*h_i_*−*h_k_*))*/*2 under induction assumption *h′_j_* = *sh_j_* and *h′_k_* = *sh_k_*, giving (*sh_j_* + *s*(*h_i_* − *h_j_*) + *sh_k_* + *s*(*h_i_* − *h_k_*))*/*2 = (*sh_i_* + *sh_i_*)*/s* = *sh_i_* = *h′_i_*). But, with dated tips, nodes closer to dated tips move less than nodes farther away from tips.

The probability of acceptance of an MCMC proposal (Green, 1995; Holder et al., 2005) is

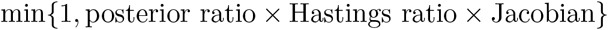

where the posterior ratio is the posterior of the proposed state *S′* divided by that of the current state *S*, the Hastings ratio the probability of moving from *S* to *S′* divided by the probability of moving back from *S′* to *S*, and the Jacobian is the determinant of the matrix of partial derivatives of the parameters in the proposed state with respect to that of the current state. The Hastings ratio has a contribution of 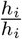 for each node that is moved, so the Hastings ratio works out as 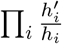. By using a Bactrian kernel, the Jacobian is 1. Note that down stretching can lead to increased branch lengths, and up stretching to reduced branch lengths, for example in the internal branch below the left branch below the root marked with dots on the nodes in Fig.1b and c respectively. In Fig.1c the dots overlap due to the the branch length being reduced to close to zero. While it is still possible for node heights to be proposed that result in negative branch lengths, if this happens often, automatic tuning parameter optimisation ensures that boldness of the move is reduced, and still a good number of proposals will be accepted.

#### 2.2.2 New epoch flex proposal

The epoch flex-operator randomly selects a lower bound *L* and upper bound *U* in the range between the root height of the tree and the youngest leaf (enforcing *L < U* by swapping values if *L > U*), then scales the interval with a random scale value *s* drawn from a Bactrian distribution (Yang and Rodríguez, 2013; Thawornwattana et al., 2018) with respect to the lower bound. Internal nodes above the upper bound *U* are moved to accommodate the scaled height of the interval. Internal nodes below *L* and leaf nodes do not have their heights changed, which allows caching of the partial calculations for the tree likelihood for at least the nodes below *L* (Fig. 2), making it a more time efficient operator that the tree stretch operator.

**Figure 2:**
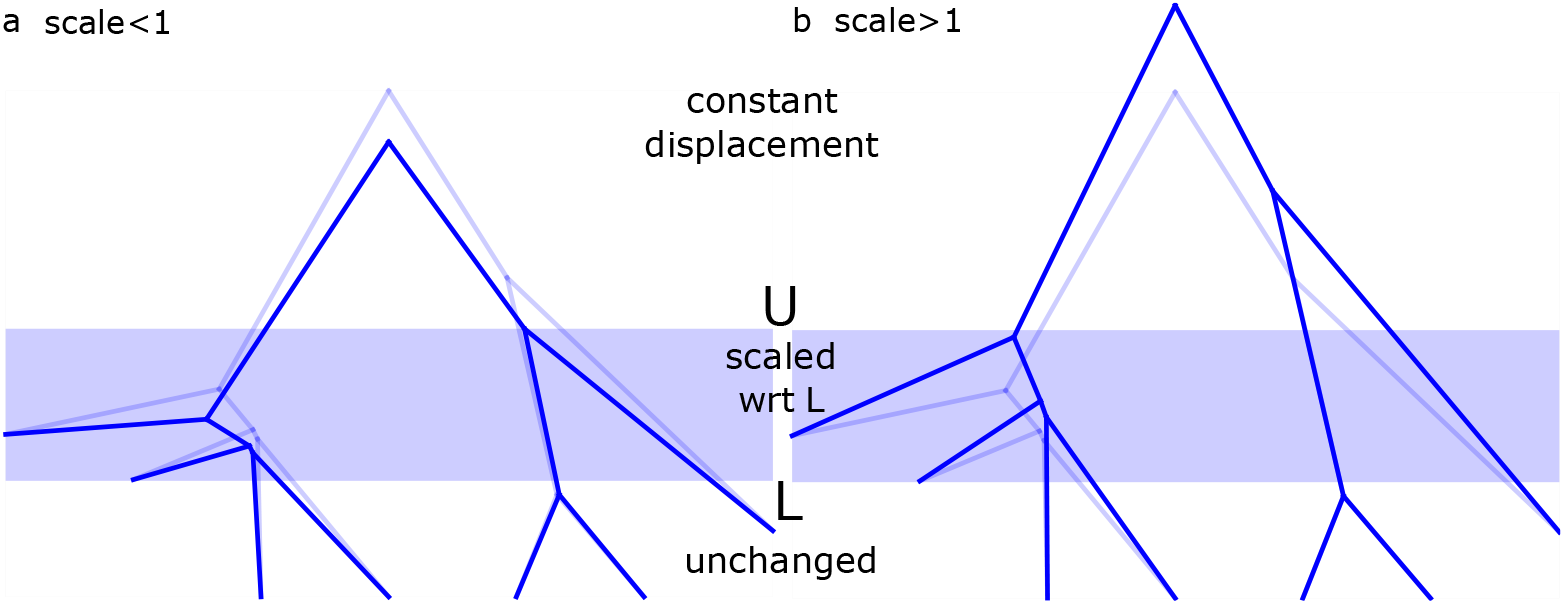
Epoch operator selects lower bound *L*, upper bound *L* and scale factor *s* and scale all nodes between *L* and *U*. Nodes above *U* are moved to make space for the newly scaled epoch. a) applied to light tree giving dark tree when scale factor less than one, and b) when scale factor larger than one.

More formally, for every node *i* with height *h′_i_* the proposed height *hi* is

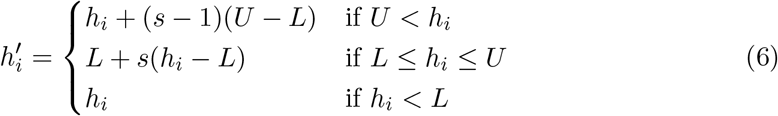

The Hastings ratio requires taking into account selecting *L* and *U* and since these are chosen uniform in the interval [0, *h_root_*] and we have a new root height 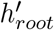 after the proposal the contribution is 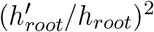 for these two random values. Furthermore, let there be *k* nodes with heights in between *L* and *U*, then the contribution of scaling these *k* nodes is *s^k^*, making the log Hastings ratio 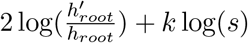.

Like for the tree stretch operator, a tuning parameter *c* is used for sampling *s* to obtain an optimal acceptance probability of 0.4. The proposal can result in direct rejection if any of the scaled nodes are assigned heights below a tip. One way to prevent this from happening is to enforce the lower bound to be older than the oldest tip, so only part of the tree above the oldest tip is scaled. Since that part of the tree tends to be less constrained by tips, bolder proposals are possible, so having both the restricted and unrestricted version of the operator in the mix can lead to better proposals overall. Note that this is only an effective strategy if there are a sufficiently large number of internal nodes above the oldest tip. This is not always the case, for example, influenza data sets can be sampled over a large duration of an outbreak, and most internal nodes may end up younger than the oldest sample.

#### 2.2.3 New up/down proposal

Mean clock rate, tree prior parameters and tree height tend to be highly correlated, so moving them at the same time (but in opposite direction) can help mixing. The so called up/down operator in BEAST randomly picks a scale factor *s* and scales up the tree with factor *s* while scaling down the clock by scaling with factor 1*/s*. Tree prior parameters like birth rate or population size can be scaled in the appropriate direction at the same time.

The new tree stretch and epoch flex operators also change tree height, so we can use *s* = *h_root′_ /h_root_* as scale factor in a similar fashion as for the up/down operator and scale clock rates and tree prior parameters. For each scaled parameter, a contribution of *s* when scaling up (or 1*/s* when scaling down) must be added to the Hasting ratio.

## 3 Validation

We performed a well calibrated simulation study in order to make sure our implementation is correct, and performed an analysis of SARS-CoV-2 for community outbreaks in New Zealand.

### 3.1 The implementation is correct

To establish correct implementation of BICEPS, we performed a well calibrated simulation study sampling 50 tip dates randomly from the interval 0 to 1. To establish correctness of the new operators, we use a coalescent tree prior with constant population size (log-normal(*μ* = 1, *σ* = 1.25) distributed), a HKY model with kappa log-normal(*μ* = 1, *σ* = 1.25) distributed and gamma rate heterogeneity with four categories with shape parameter exponentially distributed with mean=1, and frequencies Dirichlet(1,1,1,1) distributed. Further, gamma is lower bounded by 0.1 to give reasonable range of rates (Bouckaert, 2020) and frequencies lower bounded by 0.2 to prevent atypical parameter values. We use a strict clock where the clock rate times tree height has a tight normal(*μ* = 1, *σ* = 0.05) prior. Sampling 100 instances from this distribution using MCMC in BEAST 2 (Bouckaert et al., 2019), we get a range of tree heights from 1.03 to 8.8 with mean 1.6 (note that due to the tips being sampled from 0 to 1, the tree height is lower bounded by 1) and a clock rate range of 0.1 to a fraction over 1 in our study. With these trees, we sample sequences of 1000 sites using the sequence generator in BEAST 2.

Tables 1 and 2 show the coverage of true parameter values (and some other statistics) used to simulate the sequence data by the 95% highest probability density (HPD) intervals estimated after running MCMC. With 100 experiments, the 95% HPD of the binomial distribution with p=0.95 ranges from 91 to 99 inclusive. All analyses were run for 20 million samples, which was sufficient to obtain effective sample sizes of at least 200 for each of the parameters shown in Tables 1 and 2. All coverages observed are in the expected range, suggesting no problems with the implementation.

**Table 1:** Coverage of the true value by 95% HPD estimates from 100 independent runs of BICEPS for various parameters in the model and for different operators added to the standard set of operators. All coverage is in the expected 91 to 99 range, providing confidence there are not errors in the operator implementation.

| Parameter | epoch flexer | tree stretcher | up/down |
| --- | --- | --- | --- |
| tree height | 91 | 95 | 94 |
| tree length | 94 | 91 | 92 |
| kappa | 96 | 99 | 99 |
| gamma shape | 99 | 98 | 96 |
| population parameter | 97 | 94 | 93 |
| clock rate | 96 | 99 | 98 |
| tree prior | 98 | 92 | 93 |
| frequencies A | 95 | 91 | 92 |
| frequencies C | 97 | 94 | 95 |
| frequencies G | 95 | 93 | 95 |
| frequencies T | 96 | 96 | 96 |

**Table 2:** Results for 100 BICEPS analysis with 50 taxa, 250 sites and unlinked and linked population sizes with standard operators and with the new operators added in. Coverage as in Table 1 for unlinked BICEPS with standard operators, linked BICEPS with and without new operators. Effective sample size (ESS) shown compares the standard with new operators, where bold numbers indicate the better ESS. Coverage is in the expected 91 to 99 range for all cases but ESS increase for tree related parameters, in particular the minimum ESS of the 100 runs increases significantly. Coverage of the true value by 95% HPD estimates from 100 independent runs of BICEPS for various parameters in the model and for different operators added to the standard set of operators.

| Parameter | Coverage |  |  | Average ESS |  | Minimum ESS |  |
| --- | --- | --- | --- | --- | --- | --- | --- |
|  | unlinked | linked<br>standard | new | linked<br>standard | new | linked<br>standard | new |
| tree height | 97 | 93 | 94 | 1640 | **1709** | 116 | **1219** |
| tree length | 99 | 97 | 94 | 1540 | **1669** | 113 | **1200** |
| population size | 94 | 97 | 96 | 1709 | **1736** | 783 | **1250** |
| coalescent prior | 98 | 98 | 94 | 1547 | **1690** | 121 | **1216** |
| pop size epoch 1 | 97 | 96 | 98 | 1721 | **1731** | 969 | **1358** |
| pop size epoch 2 | 97 | 96 | 93 | 1704 | **1728** | 250 | **1401** |
| pop size epoch 3 | 99 | 96 | 95 | 1711 | **1731** | 232 | **1285** |
| pop size epoch 4 | 97 | 96 | 97 | 1695 | **1711** | 364 | **1343** |
| pop size epoch 5 | 94 | 95 | 96 | 1693 | **1716** | 289 | **1035** |
| kappa | 97 | 96 | 93 | 1530 | **1536** | 1114 | **1132** |
| gamma shape | 92 | 99 | 93 | 1532 | **1543** | 644 | **741** |
| frequencies A | 91 | 95 | 97 | **790** | 778 | 213 | **342** |
| frequencies C | 94 | 96 | 94 | 750 | **752** | **390** | 349 |
| frequencies G | 91 | 93 | 98 | **787** | 760 | **425** | 369 |
| frequencies T | 92 | 89 | 95 | **789** | 761 | **293** | 228 |
| clock rate | 96 | 92 | 94 | 1622 | **1690** | 125 | **1182** |

### 3.2 COVID-19 in New Zealand

We use the 887 full genome sequence data from Douglas et al. (2021a) containing samples from the 11 community outbreaks in New Zealand plus closely related sequences from the rest of the world. Further, we use a subsample of all taxa sampled up to 31 August 2020 consisting of 257 taxa for performance comparison. The data was analysed as follows. Genomic sites were partitioned into the three codon positions, plus non-coding, as described by (Douglas et al., 2021b). For each partition we model evolution with an HKY substitution model with log-normal(*μ* = 1, *σ* = 1.25) prior on kappa, frequencies estimated with Dirichlet(1,1,1,1) prior, and relative substitution rates with Dirichlet(1,1,1,1). We use a strict clock model with log-normal(*μ* = −7, *σ* = 1.25) prior on mean clock rate as in Douglas et al. (2021b,a), and for tree prior we use a Bayesian skyline model (Drummond et al., 2005) with Markov chain distribution on population sizes and log-normal(*μ* = 0, *σ* = 2) on first population size and compare this with a BICEPS tree prior. MCMC analyses were initialised with a neighbour joining tree.

## 4 Results

### 4.1 Operator performance analysis

Fig. 3a-c show violin plots for effective sample sizes (ESS) obtained with the 100 runs for the posterior, prior and tree length where the first item was done with standard operators, the second with the epoch flex operator added and the the third with tree stretch operator added as well. There was some beneficial effect from these operators on the posterior, more so on the prior, as well as the tree length. Site model parameters were practically unaffected by adding these operators, but there was some beneficial effect on the ESS for the clock rate. Note these ESSs were obtained under similar run times, so the plots suggest the operators are moderately beneficial for data simulated under the model, or at least no detrimental to mixing. However, for empirical data we observed more marked differences (see below).

**Figure 3:**
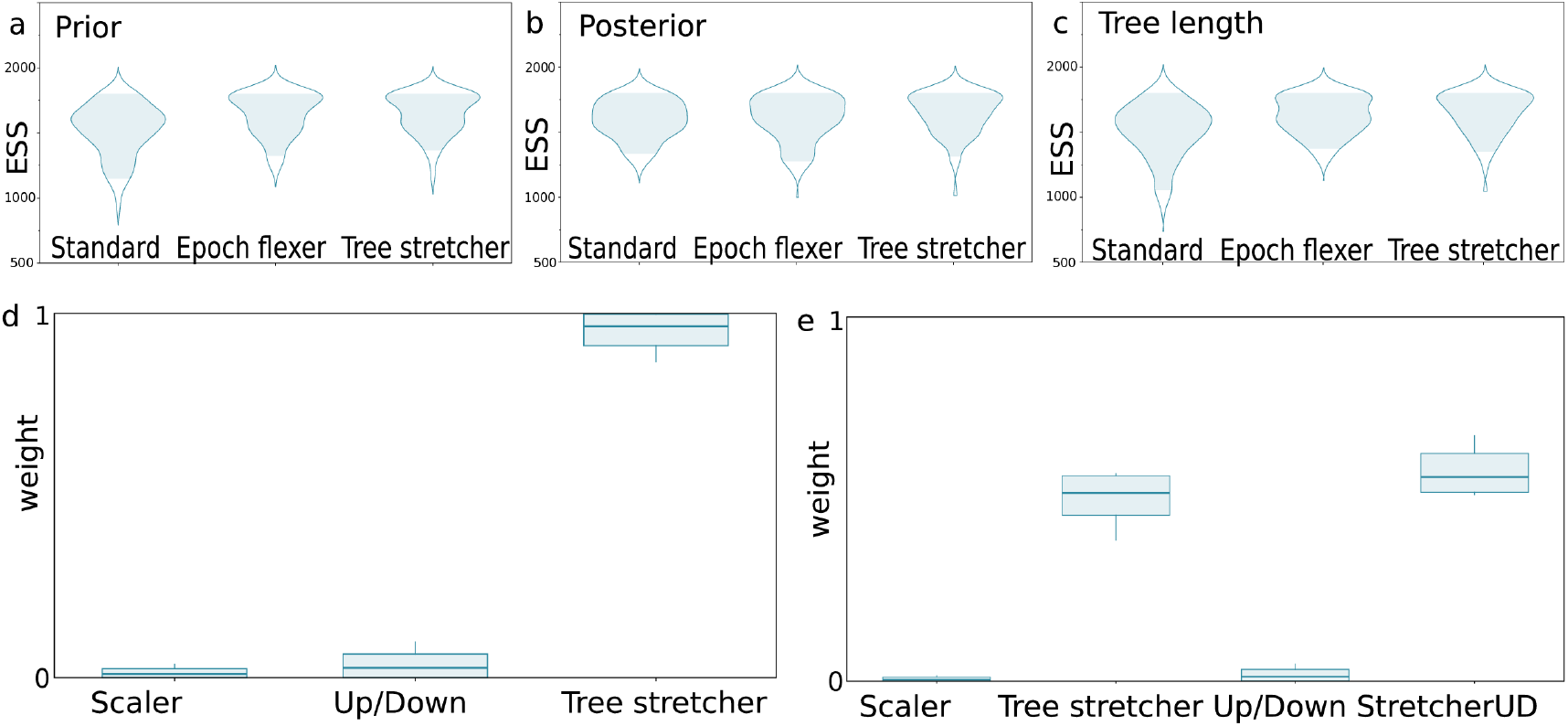
Performance of epoch flexer and tree stretch operators. Operator are weighted such that run times of various combinations are similar, so ESSs are comparable. ESSs improve a little, and weights favour the new operators over standard ones. a) ESSs on a scale of 500 to 2500 of the prior for different operator combinations – classic, with epoch operator and with epoch operator and scaler. b) ESSs for posterior, c) ESSs for tree length. d) Weights on a scale of 0 to 1 assigned to operators by the adaptable operator sampler for standard tree scaler, up/down operator and tree stretcher. e) Weights for standard tree scaler, tree stretcher, up/down operator and tree stretcher with up/down combination.

Another way to get a sense of the performance of the BICEPS operators compared to standard operators is by employing the adaptable operator sampler (Douglas et al., 2021c). This is an operator that selects among a set of operators by keeping track of relevant performance indicators of the various operators, namely amount of change in node heights, amount of time required to calculate the new state, and probability of acceptance. Together, these factors are used by the adaptable operator sampler to re-weigh sets of operators for optimal amount of node height change per unit of time.

Fig. 3d,e show the end weight distribution over the 100 runs of the well calibrated simulation study for the case where the tree scaler, up/down operator and tree stretcher were reweighted by an adaptable operator sampler, and Fig. 3e the case where a new up/down operator was added. In the first case shown in Fig. 3d, an overwhelming amount of weight is distributed towards the tree stretcher. In the second case shown in Fig. 3e, about standard operators hardly get any weight assigned, while most of the weight is distributed almost evenly between tree stretcher and new up/down operator, with a slight preference for the up/down operator. This illustrates the new tree stretch and up/down variant perform well when balancing the size of change, the time to recalculate the posterior and how often the operator is accepted. Since there is no directly comparable version of the epoch flexer, we omitted it from the mix.

### 4.2 COVID-19 analysis

For the 10 runs of the 257 taxa SARS-CoV-2 analysis, MCMC convergences (all parameters having ESSs larger than 200) around 20 million samples for the BICEPS analysis while the BSP analysis still struggles to achieve mixing. In particular the tree length only achieves single digit ESSs or ESSs less than 20 when taking favourable burn-in values for the 10 runs in Tracer. Fig. 4a shows a typical trace of the tree length for one of the BSP and one of the BICEPS analyses, highlighting how BICEPS achieves convergence much faster. Fig. 4b displays the poster ESSs over 10 runs, and shows that adding the BICEPS operators helps mixing with the BSP model. Further, integrating out parameters as done in the BICEPS model improves ESSs a bit more and fixing group sizes instead of estimating them improves ESSs even more.

**Figure 4:**
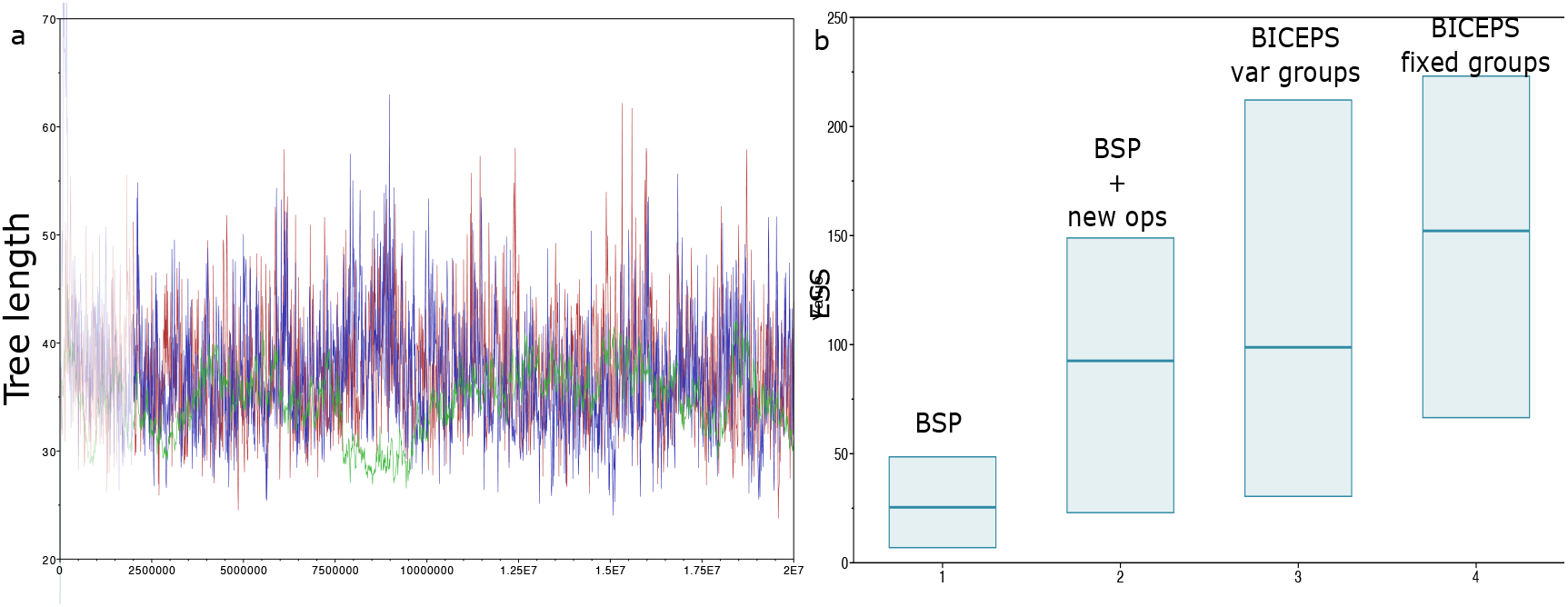
MCMC efficiency a) Trace of tree length for BSP (green line with very low period), BSP with new operators (blue line with higher period) and BICEPS analysis (red line forming a satisfying hairy caterpillar pattern). The BSP analysis typically does not reach an ESS of 10 when the BICEPS analysis already has ESSs around 200. b) Posterior ESS for 10 runs of BSP, BSP + new operators, and BICEPS with variable and fixed group sizes. Both new operators and the BICEPS prior contribute to improving ESSs.

The COVID-19 analysis from Douglas et al. (2021a) required 8 chains running 1 billion samples each and were combined to obtain satisfactory ESSs over 200. In contrast, for the same analysis with BICEPS prior and new operators a single run converged in 1 billion samples to ESSs over 200, a factor 8 speed up. Since these analyses use a different though related tree prior, we compare the tree posteriors (see Fig. 5a) and conclude these tree priors lead to very similar results in posterior tree distributions. Clade support is very similar (red dots in Fig. 5a) except for a handful of clades, which may be due to imperfect mixing of the trees, something not unexpected with this many taxa and sequences with relatively little variation (some sequences are even identical).

**Figure 5:**
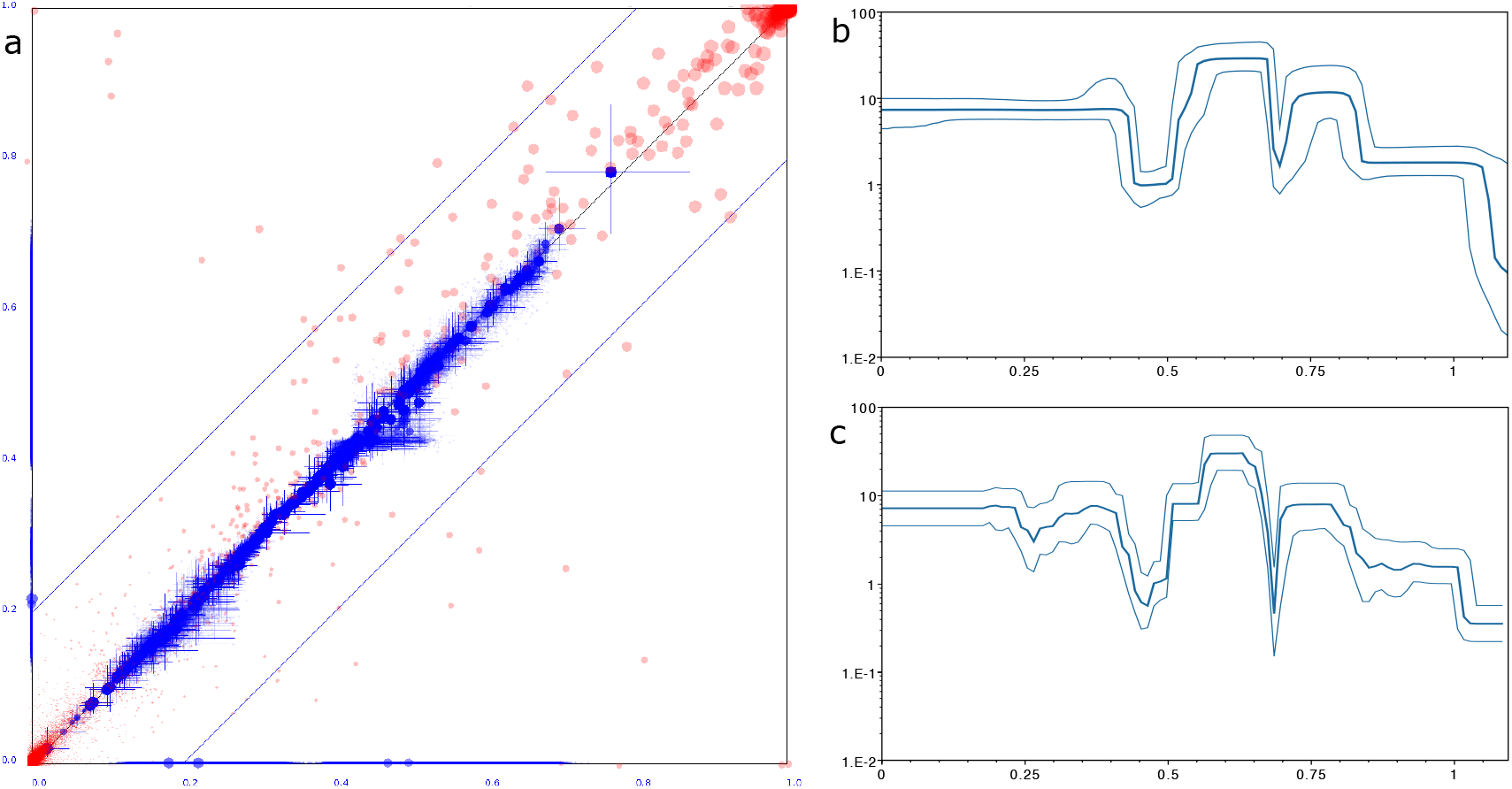
BSP and BICEPS compared. a) Difference in clade support and clade heights for the Bayesian skyline analysis from Douglas et al. (2021a) and the same analysis with a BICEPS prior. Red dots indicate clade support between 0 and 1 on both axis, blue dots indicate mean clade heights with cross hairs showing the 95% HPD intervals of height estimates. The axis are scaled between zero and the highest tree height found in either tree set. b) Population history for SARS-CoV-2 inferred with the BSP model and c) BICEPS model. Dark middle line indicates the median, lighter outer lines cover the 95% HPD intervals. The x-axis shows time in years going backward from left to right, the y-axis shows population size on a log scale. BSP and BICEPS analyses largely agree.

The estimate of most clade ages and in particular the root ages are consistent with each other. However, the BSP analysis puts the root age a fraction lower (at 1.24 year) than the BICEPS analysis (at 1.25). This can be explained when considering the demographic reconstruction, shown for BSP in Fig. 5b, and for BICEPS in Fig. 5c. Over all, the reconstructions are quite similar, but note that the population size estimates near the root (right hand side of plots) are lower and with higher uncertainty for the BSP reconstruction. The BICEPS reconstruction assumes a constant population size for each epoch and number of coalescent intervals are fixed to 29 or 30 (making 30 groups for 887 taxa). Therefore, the last 29 coalescent events to the root are assumed to be under a constant population. The BSP analysis on the other hand estimates group sizes and it is 22 on average with 95% credible range of 11 to 35, resulting in a smaller population size estimate, hence a slightly reduced root age estimate. When running the BICEPS with 10 epochs instead of 30, the effect is enlarged (giving a root age estimate of 1.29 year).

A general rule of thumb in statistics is that 30 observations are sufficient to estimate the mean of a parameter. Given that epochs can be linked through posterior mean population size estimates in BICEPS using epochs that cover more than 30 observations does not seem necessary. By default, the model uses 10 groups unless group sizes are larger than 30, then the group count is set to the number of taxa divided by 30. However, if group sizes are less than 6 then group count is set to the number of taxa divided by 6.

### 4.3 HCV analysis

To demonstrate BICEPS does not only perform well with serially sampled data, we analysed a dataset of 63 hepatitis-C virus sequences sampled in Egypt in 1993, which was earlier analysed in Drummond et al. (2005) and Stadler et al. (2013). We analysed with a GTR substitution model with gamma rate heterogeneity with 4 categories and fixed the clock rate at 7.9 × 10^−4^ substitutions per site per year.

Where BSP requires 30 million samples for MCMC to converge, BICEPS requires only 5 million samples, demonstrating that BICEPS can be considerably faster. A comparison similar to shown in Fig. 5 for SARS-CoV-2 can be found in Appendix A. It demonstrates that the BSP and BICEPS models result in very similar tree sets, but the BICEPS analysis can be performed more efficiently, both when tips are sampled through time as in the case of the SARS-CoV-2 data, or when tips are sampled at the same time as for the HCV data. This suggests that we can analyse larger data sets using the BICEPS model than the BSP model. So, the primary benefit of using this model is being able to analyse more sequences and allowing us to investigate processes such as demographic reconstructions in more refined detail.

### 4.4 Generalisation to other tree priors

The efficiency of the BICEPS tree prior relies on integrating out population sizes, so that fewer parameters need to be inferred. Here, we used an inverse gamma distribution over population sizes, but a gamma distribution would be a suitable alternative. For models with more parameters, like the besp tree prior which takes sampling in account Parag et al. (2020), integrating out parameters analytically if possible at all would require non-standard techniques. Regardless, coalescent models assume that the samples represent a small number of individuals from a much larger population. When this assumption does not hold, birth death models may be more appropriate. However, it is more challenging to extend the idea of integrating out parameters to birth death sampling models.

For the Yule model (Yule, 1924; Aldous, 2001), a pure birth model, this is straight-forward (Appendix B). An epoch version of the Yule model assuming death and sample rates of zero and sampling all extant taxa at the same time (that is, rho-sampling with rho=1) can be found in Appendix C. The latter is available as ‘Yule skyline’ model in BEAST in the BICEPS package. This provides a flexible prior for the case where tips are not sampled through time, but are all taken at the same time. The model is implemented in BEAST 2 and a well calibrated simulation study (Appendix C) passed. For more general cases this approach is hampered by the large number of parameters (birth, death, sampling rate, etc.) and because the tree likelihood is of a form that does not appear to lend itself for integrating out parameters.

The BICEPS and Yule skyline tree priors put coalescent events in approximately equally sized groups in order to reduce noise and provide estimates of population sizes and birth rates respectively with tight uncertainty bounds. An alternative is to split the tree height into equally sized time intervals and use the coalescent and lineage count information in these same sized epochs. Though most epoch boundaries do not coincide with coalescent events any more, this has little impact in the way the mathematics works out, but will impact the distribution of coalescent events in the intervals: usually, there will be fewer near the root and more near sampling times. Consequently, uncertainty bounds will become larger near the root and smaller in epochs containing larger numbers of coalescent events.

### 4.5 Primates analysis

A primate alignment of full mitochondrial genomes with 87 taxa and 19220 sites (Finstermeier et al., 2013) was analysed using a GTR substitution model with estimated frequencies, optimised relaxed clock model (Douglas et al., 2021c) and Yule tree prior (see supplementary material for BEAST 2 XML files for this and associated analyses). Due to the very informative sequence data, this analysis tends to mix slowly because the posterior is very peaked making it hard for standard operators to make bold moves (Zhang and Drummond, 2020). Fig. 6 shows how adding the BICEPS operators does help mixing of the posterior and the likelihood, demonstrating that adding BICEPS operators allows analyses to run more efficiently. A Yule skyline analysis with the same data shows significant improvements in mixing for both posterior and likelihood compared to the Yule analyses with standard operators, but slight degradation of the posterior ESS though still improved likelihood ESS compared to Yule analyses with BICEPS operators. A birth rate skyline reconstruction through time shows that there is only small variation through time. In fact, Fig. 6c shows the mean birth rate under the Yule model, which assumes a constant birth rate throughout the whole tree, as dashed horizontal line. The line fits inside the whole 95% HPD trajectory, which suggests a constant rate of speciation of primates can not be ruled out.

**Figure 6:**
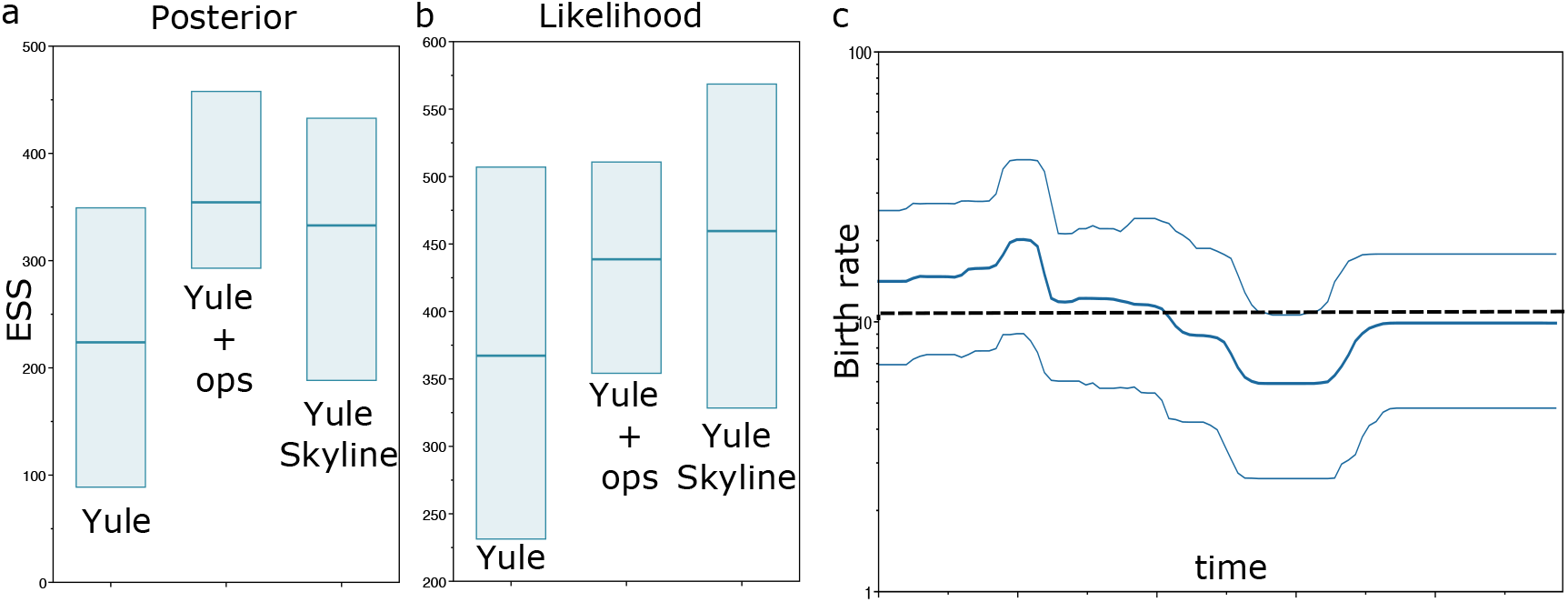
Primate analysis. a) ESS over 10 runs for posterior when using Yule with standard operators, Yule with BICEPS operators and Yule skyline with BICEPS operators. b) ESSs for the likelihood. Note the change in scale. c) Reconstruction of birth rates with Yule Skyline showing median and 95% HPD intervals. The dashed line shows the mean birth rate for a Yule analysis.

## 5 Conclusions

We introduced a two headed approach for improving the efficiency of Bayesian inference under epoch models: a flexible tree prior based coalescent epoch model that integrates out population size parameters and a set of new MCMC proposals directly targeting tree lengths. Both these elements contribute to more efficient inference, in particular with SARS-CoV-2 data and with serially sampled sequence data. The behaviour of BICEPS tree prior is very similar to that of the popular Bayesian skyline plot, and allows for reconstruction of demographic histories through time making it possible to estimate timing and magnitude of population bottlenecks as well as track population expansions through time.

A generalisation to a pure birth prior under an epoch model that integrates out birth rate parameters, the Yule skyline model, is detailed in Appendix C. Other generalisations integrating out tree prior parameters appear to be mathematically challenging. The benefit of integrating out parameters instead of estimating them through MCMC as well as the more efficient tree operators is that it becomes possible to analyse larger datasets and infer more detailed population histories. Even if the population history is of no interest, but for example the tree topology, timing of origins of clades or evolutionary rate estimates are the topic of investigation, the BICEPS model provides a flexible tree prior that caters for a wide range of tree shapes and sizes with little requirements in terms of prior knowledge, unlike many birth death based priors.

The application of the new tree operators is not limited to the BICEPS tree prior, but can be used in combination with any tree prior. These operators can be expected to contribute to more efficient inference under a wide range of models, and make it possible to include more taxa than is possible with the currently available standard set of operators. This is especially important with the growing amount of sequence data, and allows for more detailed post hoc analyses by techniques such as lineage through time plots, or when location information for taxa is available, introduction through time plots (see Douglas et al. (2021b) for an example applied to COVID-19). Most tree operators in BEAST either move a very small number of nodes (often just one), or move all nodes. The tree stretch operators introduced here moves all nodes, while the epoch flex operator moves a large subset of nodes. A tree operator that randomly selects a single node, proposes a new height and moves surrounding nodes to accommodate the node height change by minimising changes in evolutionary distances did not prove to be effective in that it did not increase effective sample sizes per unit of time. It is an open question whether tree operators for Bayesian inference under MCMC that move a small subset of nodes can contribute to the efficiency of MCMC.

The BICEPS tree prior and operators are implemented in BEAST 2 Bouckaert et al. (2019) and can be used in combination with a large range of different data types, substitution and site models as well, a number of clock models, sampled ancestor trees and in combination with various types of data, including geographical locations, morphological characters, micro satellite, etc.

## Availability

The open source BICEPS package for BEAST 2 (Bouckaert et al., 2019) is available under GPL at https://github.com/rbouckaert/biceps. An analysis can be set up through BEAUti, the user friendly GUI for BEAST, both for the BICEPS and Yule Skyline models.

## Acknowledgements

I thank Alexei Drummond and Jordan Douglas for stimulating discussions, Jordan Douglas and Cinthy Jimenez-Silva for proofreading the manuscript and anonymous reviewers for providing useful comments that helped improve the manuscript.

## Supplementary material

BEAST XML files used in the experiments are available at https://github.com/rbouckaert/biceps/releases/tag/v0.0.1.

## Appendices

## Appendix A BSP vs BICEPS comparison for HCV

**Fig. S1:**
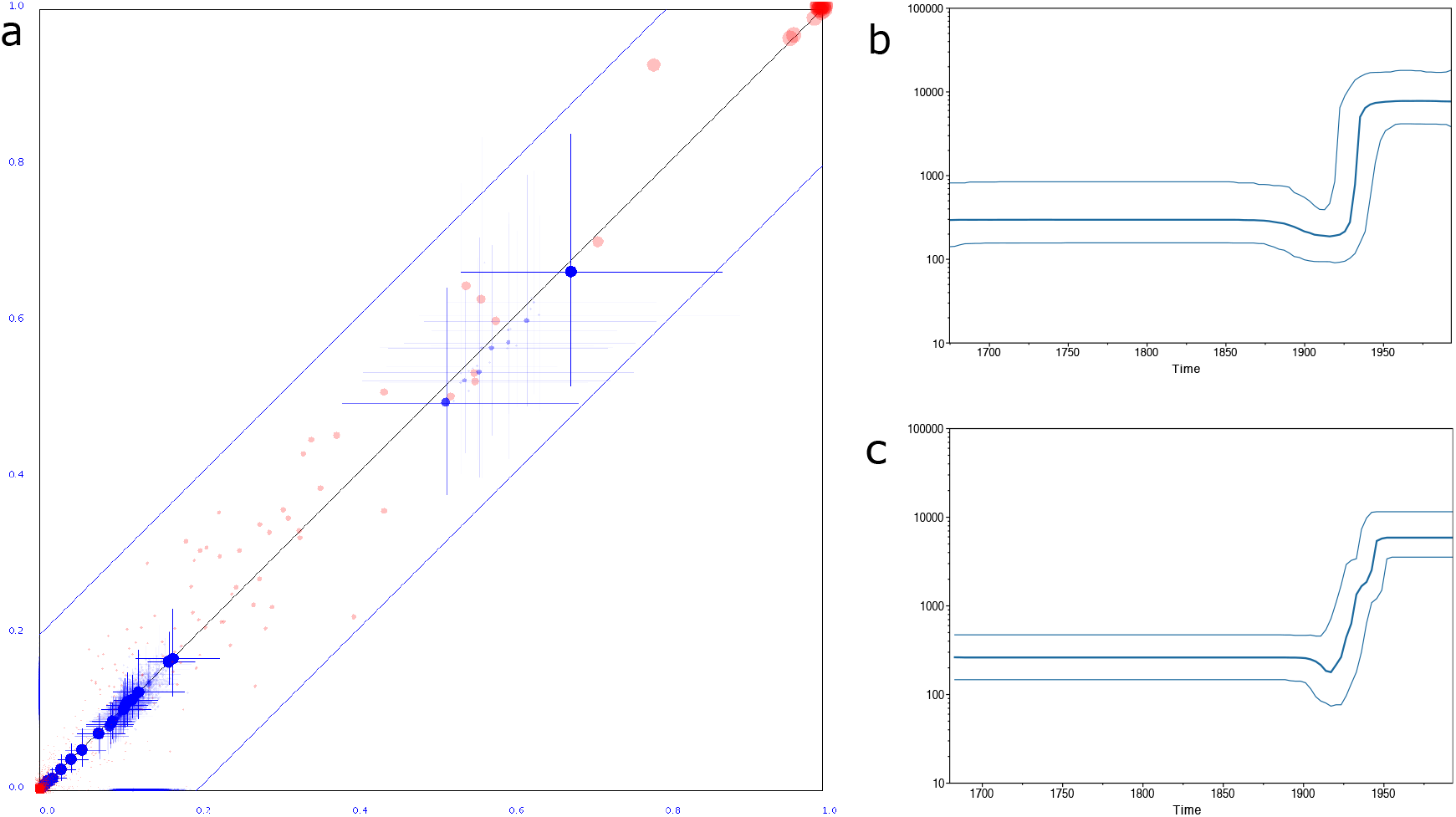
Clade set comparison as in Fig.5. a) BSP on y-axis, BICEPS on x-axis. b) BSP and c) BICEPS demographic plots

## Appendix B Yule prior

Consider a binary tree *T* with *n* taxa, leaf nodes numbered 0, *…, n* − 1, internal node numbered *n, …,* 2*n* − 2 and root numbered 2*n* − 1. Let *h_i_* denote the height of the *i*th node.

The Yule prior is defined as *pY_ule_*(*T* |*λ*) = *λ*^*n*−1^*e^−λL^* where 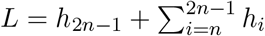 is the sum of lengths of all branches in *T*, or the length of the tree and *λ* the birth rate.

## Uniform prior

Assuming uniform prior on *λ* on [0, *u*), integrating out *λ* means solving 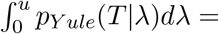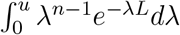, which equals *u^−^* (*Lu*)^(−(*n*−1)^ (Γ(*n*) − Γ(*n, Lu*))*/L* where Γ[*n, Lu*] the incomplete gamma function. This simplifies to

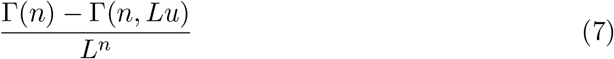

## Gamma prior

Assuming gamma prior on *λ* with shape parameter *α* and rate parameter 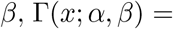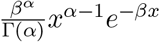. Integrating out *λ* means solving

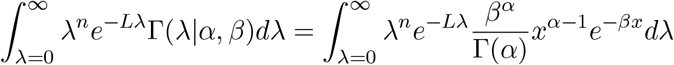

Putting constants outside the integral, and combining terms gives

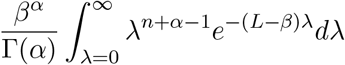

Now note that the part inside the integral is of the form Γ(*x*; *α* + *n, β* + *L*) but divided by the constant 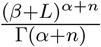 so we can write

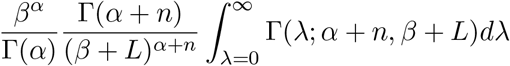

and since the gamma density integrates to 1, this leaves

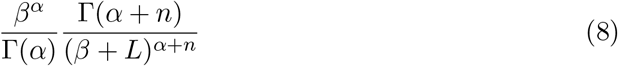

In summary, we can integrate out a uniform (Eq(7)) or gamma prior (Eq(8)) on the birth rate for a Yule prior. Note that the Yule prior is conditioned on all leafs being sampled at the same time, so this should not be used when sampling tips through time.

## Appendix C Yule skyline prior

Consider *m* epochs with assignment *A* defined as for the coalescent prior before and let Λ = {*λ*_1_, …, *λ_m_*} be a vector of *m* birth rates. We are interested in the density of a tree *T* given *A* and Λ *f* (*T* |Λ, *A*).

Starting with the general birth-death skyline model defined in Theorem 1 of Stadler et al. (2013) and assuming no sampled tips through time, setting death, sampling rates to zero for all epochs (∀*_i_μ_i_* = *ψ_i_* = 0) and only allow sampling of all lineages (*ρ* = 1) at the same time we obtain for all 0 < i *< m*, *A_i_* = *λ_i_*, *B_i_* = 1 and *q_i_* = 1*/exp*(−*λ*(*t* − *t_i_*)). Filling this into the tree likelihood in Theorem 1 of Stadler et al. (2013) gives us

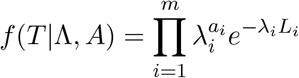

where *L_i_* the sum of the lengths of branches inside epoch *i* (remember *a_i_* is the number of coalescent events for epoch *i*). Note that each term in the product 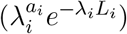 is of the same form as for the Yule prior, so we can integrate out *λ_i_* under a uniform prior bounded by 0 and finite upper bound *u*. We can also assume a gamma distribution Γ(*x*; *α, β*) for each of the epochs and obtain posterior

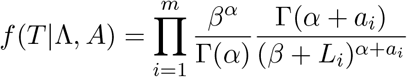

As for BICEPS, epochs can be linked obtaining a smoothing prior; we fix the shape parameter *α*, but instead of having a fixed scale *β* for each epoch, the rate *β_i−_*_1_ for epoch *i* − 1 can be chosen such that the posterior mean of the previous epoch (which is 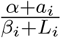 for epoch *i*) is used a priori. So, 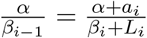 giving 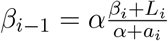

**Note 1**: The birth process moves forward in time, unlike the coalescent process, which moves backward in time. Therefore, when putting the smoothing prior on birth rates, we put a Γ(*α, β*) prior on *λ_m_*, and proceed in reverse order from epoch *m* to epoch 1 from the way the coalescent smoothing prior is calculated.

**Note 2**: Like for the Yule prior, this assumes all tips are sampled at the same time. The Yule skyline is not appropriate for data that is sampled through time.

## Validation

A well calibrated simulation study was performed with 50 contemporary taxa, and a root age prior normally distributed with mean 100 and standard deviation 0.5. The site and clock priors are as for the study for the BICEPS prior. The gamma prior used for the birth rate is with shape of 2 and rate estimated with log-normal(*μ* = 1, *σ* = 1.25), and the 8 epochs are unlinked in one set and linked in the other. Alignments of 250 sites were randomly generated. Coverage for the various parameters is listed in the table below, and it passes the test.

| parameter | coverage<br>unlinked | coverage<br>linked |
| --- | --- | --- |
| prior | 94 | 96 |
| Tree.height | 93 | 95 |
| Tree.treeLength | 94 | 95 |
| YuleSkylineBirthRateRate | 93 | 97 |
| YuleSkyline | 95 | 97 |
| BirthRates.1 | 97 | 93 |
| BirthRates.2 | 97 | 96 |
| BirthRates.3 | 94 | 92 |
| BirthRates.4 | 96 | 95 |
| BirthRates.5 | 93 | 97 |
| BirthRates.6 | 97 | 100 |
| BirthRates.7 | 95 | 95 |
| BirthRates.8 | 99 | 98 |
| gammaShape | 96 | 92 |
| kappa | 96 | 91 |
| freqParameter.1 | 92 | 95 |
| freqParameter.2 | 98 | 96 |
| freqParameter.3 | 97 | 94 |
| freqParameter.4 | 93 | 95 |
| clockRate | 93 | 99 |

## Footnotes

1 Though the notation of Drummond et al. (2005) is mostly followed here, we use *r* instead of *s* since later *s* will be used to denote scale factors for MCMC proposals.

